# scGSCA: gene-set differential co-expression analysis for single-cell RNA sequencing data

**DOI:** 10.64898/2026.10.05.756579

**Authors:** Zhuoli Jin, Rebecca Irlmeier, Jie Ping, Yuan Wang, Yan Guo, Fei Ye

## Abstract

Understanding disease-associated pathway coordination requires distinguishing phenotype-related co-expression changes from differences across cellular contexts while accounting for cells sampled from the same subject. Bulk gene-set co-expression methods do not directly address the sparse counts, single-cell distributional noise, subject-level dependence, and cell-type heterogeneity of single-cell RNA sequencing (scRNA-seq) data. We developed single-cell gene-set co-expression analysis (scGSCA), a framework for pathway-level differential co-expression analysis in scRNA-seq data. scGSCA fits gene-wise zero-inflated negative binomial mixed models with subject-level random effects and computes gene-set co-expression statistics from quantile residual correlations. It provides an overall differential gene-set co-expression analysis and an interaction test to assess whether the co-expression pattern between two cell types varies by sample phenotype. We applied scGSCA to tumor–normal gastric, breast-subtype, and lung–skin comparisons with matched bulk and scRNA-seq data. The overall test identified signals not captured in bulk analysis, including Axon guidance in gastric cancer and estrogen-response, IL6-JAK-STAT3, and TNF-*α* /NF-κB pathways in breast cancer. The interaction test further identified co-expression contrasts across cell types. scGSCA extends gene-set co-expression analysis to single-cell data while retaining subject-level structure and cell-type-specific context. scGSCA is freely available as an open-source R package.

## Introduction

Single-cell RNA sequencing (scRNA-seq) has enabled transcriptomic variation to be resolved across cell types, cellular states, and disease contexts (1–3). This resolution is particularly valuable in complex tissues such as tumors, where disease-associated transcriptional changes may be restricted to specific cell populations. Co-expression analysis can complement expression-based analyses by examining how genes vary together within biological pathways (4).

Gene co-expression analysis characterizes coordinated transcriptional variation and changes in pathway-associated expression patterns across biological conditions (5,6). In disease states, genes within a pathway may gain coordination, lose homeostatic regulation, or shift co-expression in a cell-type-specific manner. Detecting these perturbations can reveal regulatory network rewiring associated with disease progression, cellular adaptation, and treatment response (7,8). Gene-set approaches improve interpretability by evaluating predefined pathways rather than isolated gene pairs.

However, extending gene-set-level co-expression analysis to scRNA-seq data presents distinct statistical challenges. First, scRNA-seq studies often profile many cells from a smaller number of biological subjects. Cells from the same subject share genetic, microenvironmental, technical, and clinical factors, so treating them as independent observations causes pseudoreplication and inflated significance (9,10).

Second, scRNA-seq counts are often sparse and overdispersed, which can lead to unstable or biased conventional correlation estimates (11). Third, co-expression relationships are often cell-type-specific; consequently, analyses that aggregate cells across different lineages risk masking or confounding biologically meaningful signals (12). These issues compound in pathway-level inference, where gene-gene correlations are aggregated into joint test statistics and evaluated by resampling.

Several methods have been developed for differential co-expression or gene-set-level coordination analysis in bulk RNA-seq data (13–24). Among them, Gene Sets Net Correlations Analysis (GSNCA) provides a useful framework for testing whether the correlation structure of a predefined gene set differs between biological conditions by comparing the leading eigenvectors of condition-specific gene-gene correlation matrices (16). Nevertheless, these methods were developed primarily for bulk RNA-seq data and generally assume independent observations. A common extension is pseudo-bulk aggregation, which adapts bulk algorithms but obscures cell-level variation and cell-type-specific network dynamics (25).

Recent single-cell methods improve gene-gene association estimates by addressing sequencing depth, measurement error, sparsity, and local cell structure. However, most target pairwise gene-level correlations rather than formal gene-set-level testing. A recent method, GSNCASCR, represents an important step toward gene-set-level analysis of single-cell data by combining single-cell correlation estimation with the GSNCA testing framework (26). Despite these advances, a unified framework is still needed to account simultaneously for subject-level dependence, zero inflation, overdispersion, and cellular heterogeneity within formal pathway-level inference.

To overcome these limitations, we developed single-cell gene-set co-expression analysis (scGSCA), a framework for evaluating gene-set differential co-expression and sample-phenotype-dependent contrasts across cell types. Building on GSNCA, scGSCA fits zero-inflated negative binomial mixed models (ZINBMMs) with subject-level random effects and computes gene-set-level co-expression statistics from simulation-based quantile residuals (27–29). This approach accommodates overdispersion and a potential excess-zero component, while subject-level resampling preserves the sampling hierarchy.

scGSCA provides two complementary tests. The sample-phenotype test detects changes in gene-set-level co-expression structure associated with sample phenotypes, whereas the cell-type interaction test assesses whether cell-type-specific co-expression contrasts differ between sample phenotypes. We evaluated scGSCA using three case studies with bulk transcriptomic and single-cell data, spanning gastric cancer, breast cancer, and a lung versus skin comparison of normal tissues used as a biological positive control (30–34). scGSCA integrates subject-aware modeling with pathway-level co-expression analysis to evaluate sample-phenotype-associated changes and determine whether between-cell-type co-expression contrasts vary across sample phenotypes. Together, these analyses connect pathway coordination with the cellular contexts in which it differs.

## Materials and Methods

### Datasets and pathway selection

We analyzed three public datasets with matched bulk and single-cell transcriptomic data. The gastric cancer scRNA-seq dataset (Genome Sequence Archive accession HRA002336) comprises 111,140 cells from paired tumor and adjacent normal samples of 24 patients; the matched bulk RNA-seq data comprises 44 paired samples (Gene Expression Omnibus accession GSE206785). The breast cancer dataset (Gene Expression Omnibus accession GSE176078) includes bulk RNA-seq data for 11 ER-positive and 9 TNBC tumors and scRNA-seq data for 73,730 cells from 11 ER-positive and 9 TNBC patients. The GTEx data (database of Genotypes and Phenotypes accession phs000424) includes bulk RNA-seq data for 3 lung and 3 skin samples with scRNA-seq data from 40,611 cells. Cell-type annotations were taken from the original studies. For each case study, we tested a small set of biologically relevant pathways selected a priori from KEGG (gastric cancer) or the MSigDB Hallmark collection (breast cancer and GTEx). The list of genes in each pathway is provided in Supplementary Data S1.

### Bulk and pseudo-bulk comparisons

As a bulk-level reference, GSNCA was applied to the bulk RNA-seq data for the same pathways. For each cell-type pair tested in the interaction analysis, we also constructed pseudo-bulk profiles by pooling cells from the two cell types within each sample, excluding all other cell types, and applied GSNCA to the same pathway-pair combinations. Although pseudo-bulk is a common alternative, it discards cell-level variation and conflates the two cell types.

### Overview of scGSCA

The scGSCA framework is built on the GSNCA framework to test whether the co-expression structure of a predefined gene set differs between comparison groups in a single-cell setting (16). Let *G* = {1, 2, ⋯, *q*} denote an *a priori* defined gene set containing *q* genes and let *l* ∈ {1,2} denote two biological conditions. For each condition *l*, a *q* × *q* gene-gene correlation matrix 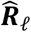 is obtained. Let *v*1_*l*_ denote the leading eigenvector of 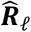 (with unit norm), representing the direction where the gene-gene correlation varies the most. The gene importance vector is defined as 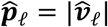, where the absolute value is taken elementwise. The difference in co-expression structure of the gene set between the two conditions is quantified using the *L*_1_ distance between the corresponding gene importance vectors

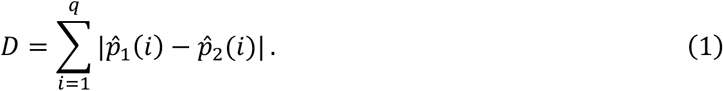

A larger value of *D* indicates a greater difference in the co-expression patterns of the gene set between the two sample phenotypes.

The original GSNCA framework assumes independent observations and was developed primarily for bulk transcriptomic data. This assumption is violated in scRNA-seq data because multiple cells are measured from the same subject and expression counts are sparse, overdispersed, and zero-inflated. Our proposed scGSCA framework retains the GSNCA test statistic *D* but modifies the correlation estimation and permutation procedures to account for these features of single-cell data.

scGSCA includes two complementary testing strategies.

1. Sample-phenotype analysis: the comparison is between two specified sample phenotypes within the given dataset. The objective is to determine whether the correlation matrices differ between the two sample phenotypes after accounting for cell-type effects and subject-level dependence.
2. Cell-type interaction analysis: the comparison determines whether the difference in correlation matrices between two cell types varies across sample phenotypes.

Sample phenotype defines the sample-level comparison groups. Samples may be from two independent groups of subjects (unpaired) or the same subjects (paired). Examples include disease conditions or treatment groups comprising different subjects and paired tumor and adjacent normal tissues from the same subjects.

Figure 1 provides a general overview of the statistical workflow of the scGSCA framework. To determine the statistical significance of both sample-phenotype and cell-type interaction analyses, we employ a non-parametric permutation framework. This procedure evaluates the significance of the observed test statistics (*D*_*obs*_ and *δ*_*obs*_) using resampling that respects the subject-level sampling structure, subject to the exchangeability assumptions described below.

**Figure 1.**
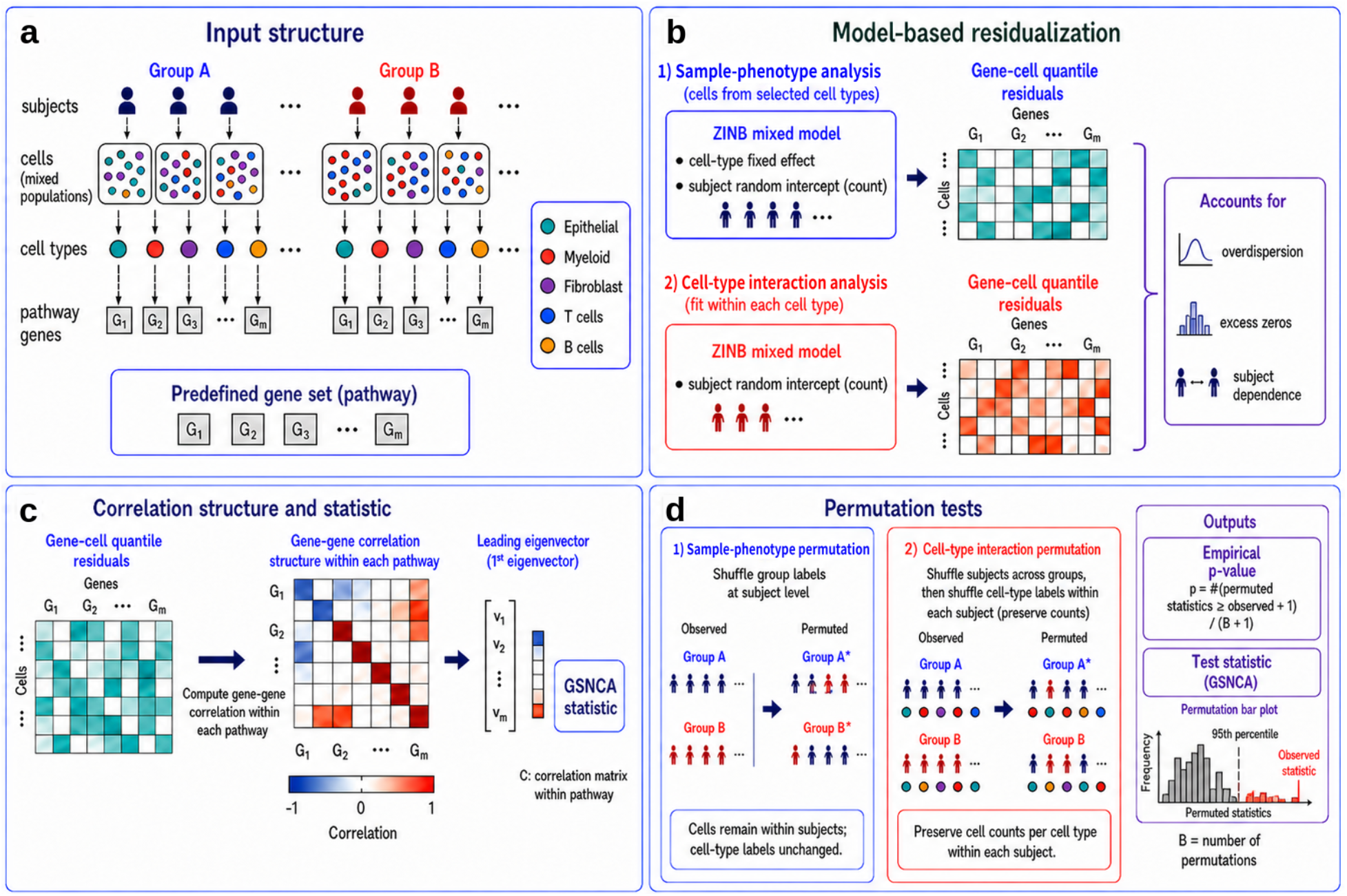
Statistical workflow of the scGSCA framework. (a) scGSCA analyzes gene sets from scRNA-seq data with nested subject, cell, and cell-type structure across two sample phenotypes. (b) Gene expression counts are modeled using a ZINBMM to account for overdispersion, excess zeros, and subject-level dependence. The sample-phenotype analysis includes cell type as a fixed effect; the cell-type interaction analysis fits models within each cell type. (c) Gene-wise quantile residuals are used to construct gene-gene correlation structures within each gene set, from which GSNCA statistics are computed. (d) Statistical significance is assessed by permutation tests that preserve subject-level structure. Alt text: Four-panel schematic of the scGSCA workflow. Panel (a) shows subjects in two groups, each contributing cells of several cell types, and a predefined gene set. Panel (b) shows zero-inflated negative binomial mixed models producing cell-by-gene quantile residual matrices for the sample-phenotype and cell-type interaction analyses. Panel (c) shows a gene-gene correlation heatmap, its leading eigenvector and the GSNCA statistic. Panel (d) shows subject-level and within-subject cell-type label permutations and a permutation null distribution with the observed statistic.

### ZINBMM

To account for overdispersion and excess zeros in scRNA-seq counts, scGSCA models gene expression using a ZINBMM. For each gene *g* ∈ *G*, let *Y*_*cgs*)_ denote the count for gene *g* in cell *c* of cell type *t*_*cs*)_ from subject *s*. We assume

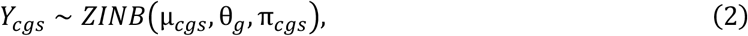

where µ_*cgs*_ denotes the mean count for gene *g* in cell *c* from subject *s*, θ_*g*_ is the gene-specific dispersion parameter, and π_*cgs*)_ is the zero-inflation probability.

Models are fitted separately for each sample phenotype. For the sample-phenotype analysis, models are fitted separately for each phenotype using all analyzed cell types. For the cell-type interaction analysis, a separate model is fitted for each phenotype–cell-type combination.

For the sample-phenotype analysis, cell type is included as a fixed effect in both the count and zero-inflation components, and subject-level dependence is modeled by a random intercept in the count component. Specifically,

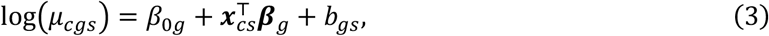

where 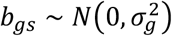, and

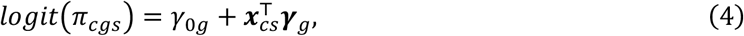

where ***x***_*cs*)_ contains indicators for non-reference cell types, *β*_,0*g*_ and *γ*_0*g*_ are gene-specific intercepts, ***β***_*g*_ and ***γ***_*g*_ are vectors of cell-type effects, and *b*_*gs*)_ is the subject-specific random intercept. This specification adjusts for cell-type heterogeneity while accounting for correlation among cells from the same subject.

Because residuals from all analyzed cell types are pooled, the sample-phenotype statistic reflects the overall co-expression structure of the cell population, including contributions from differences in cell-type composition between phenotypes.

Cell-type-specific changes are assessed by the cell-type interaction analysis. Each fitted model contains cells from only one cell type, thus no cell-type fixed effect is included. Subject-level dependence is modeled by a random intercept in the count component. Specifically,

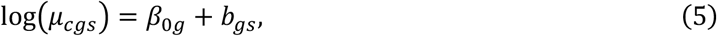

where 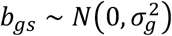, and

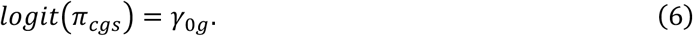

This specification accounts for correlation among cells from the same subject while estimating gene-level residual variation within each cell type. In both analyses, we fit the models using a custom C++ implementation that follows the zero-inflated negative binomial mixed-model formulation of glmmTMB, and maximizes a Laplace-approximated marginal likelihood with NLopt (version 2.10.1) (27,35–37). For each permutation, we refit the models using the permuted labels and recalculate the quantile residuals because the models are not permutation invariant.

### Quantile residuals

Pearson correlation is valid for bulk data, but its application to single-cell data must address sparsity, overdispersion, and measurement noise. We therefore standardize residuals from the fitted ZINBMM to obtain quantile residuals for correlation estimation.

After fitting the ZINBMM for gene *g*, let *y*_*cgs*)_ denote the observed count of gene *g* in cell *c* from subject *s*, and let 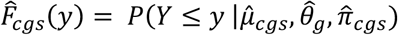 denote the fitted ZINB cumulative distribution function (CDF).

Following Dunn and Smyth (28), we draw a uniformly distributed random variable

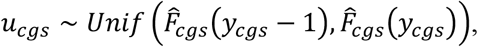

and the quantile residual is defined as

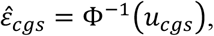

where Φ^.−1^ denotes the inverse CDF of the standard normal distribution. Under a correctly specified model, the quantile residuals are approximately standard normal. In practice, we compute these simulation-based quantile residuals using the DHARMa package (29). DHARMa uniform residuals were transformed using the standard normal quantile function before correlation estimation.

Specifically, for each comparison group (sample phenotype or cell type), we form the *n* × *q* quantile residual matrix 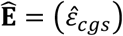. Here, *n* is the total number of cells in each group, and *q* is the number of genes. The *q* × *q* gene-gene correlation matrix 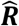 is computed as the matrix of pairwise absolute Pearson correlations among the standardized columns of Ê (denoted as 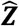), i.e.

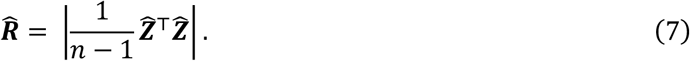

These correlation matrices then serve as input to the GSNCA test statistic *D*. The resulting associations describe dependence between model-based residuals.

### Permutation test

Statistical significance is assessed using non-parametric permutation procedures tailored to scRNA-seq data. To preserve within-subject dependence and the hierarchical structure of the data, all cells belonging to a subject or paired specimen receive the same reassigned sample-phenotype label. Owing to runtime, we used 200 permutations for single-cell sample-phenotype tests and 500 permutations for cell-type interaction and GSNCA tests.

For sample-phenotype analysis, we test the equivalence of the gene importance vectors (absolute leading eigenvectors) between two phenotypes: *H*,: ***p***_1_ = ***p***_2_ *vs. H*_1_: ***p***_1_ ≠ ***p***_2_. The *P* value of this test is evaluated through permutation. Let *n*_*perm*_ be the number of permutations, *D*_*obs*_ be the observed test statistic, and *D*_*perm,i*_ be the test statistic of the *i*th permutation, then

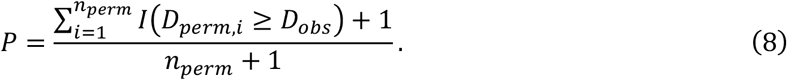

Group labels are permuted (only) at the subject level. For paired designs, sample-phenotype labels are swapped with probability 0.5 independently across subjects. For unpaired designs, such as subtype or tissue-specific comparisons, sample-phenotype labels are randomly reassigned at the subject level while preserving the original sample-phenotype proportions. For each valid permutation, ZINBMMs are refitted, quantile residuals are recalculated, residual-based correlation matrices are reconstructed, and the permuted test statistic *D*_*m*_ is computed for the *m*^th^ permutation.

For cell-type interaction analysis, we compute the difference of *D* for each sample phenotype. Denote 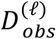 as the observed statistic *D* for sample phenotype *l* (*l* = 1,2), and 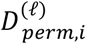 as the permuted statistic *D* for sample phenotype *l* of the *i*th permutation. The final statistic is 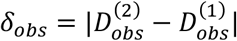, representing the magnitude of the difference in between-cell-type co-expression contrasts across sample phenotypes. The permutation statistic *δ*_*perm,i*_ and *P* value are defined analogously. Under the null hypothesis, conditional on subject membership and the number of cells of each type within each subject, the joint distribution of the data used to construct the cell-type interaction statistic is assumed to be invariant to permutations of the cell-type labels within subjects. We additionally assume the cell-type labels are exchangeable within subjects under the null hypothesis. Accordingly, for the interaction test, subject-level phenotype permutation is followed by cell-type label shuffling within each subject, preserving cell-type counts. ZINBMMs are refitted, and quantile residuals, correlation matrices, and the cell-type interaction test statistic are recomputed for each valid permutation.

### Implementation

The proposed scGSCA framework was implemented as an R package, with performance-critical components written in C++ via Rcpp, for gene-set-level differential co-expression analysis of scRNA-seq data (38).

## Results

Table 1 compares scGSCA with existing gene-set and single-cell co-expression methods across key analytical and statistical features. Bulk RNA-seq methods such as GSCA, GSDCA, GSNCA, MetaGSCA, EDDY, KEDDY, and CoGA provide gene-set-level differential co-expression testing, whereas single-cell co-expression methods such as COTAN, CS-CORE, and GSNCASCR address single-cell data or cell-type-specific associations. However, none jointly provide cell-type-aware inference, hierarchical dependence, count-aware modeling, and sparsity within gene-set-level inference. scGSCA was developed to bridge this gap by combining gene-set co-expression statistics, ZINBMM-based residualization, and cell-type-aware permutation testing (Table 1).

**Table 1.** Comparison of scGSCA with existing gene-set co-expression and single-cell co-expression methods.

| Method | RNA-seq data type | Publication year | Gene-set-level statistic | Cell-type-aware | Hierarchical-structure-aware | Sparsity-aware |
| --- | --- | --- | --- | --- | --- | --- |
| GSCA | Bulk | 2009 | ✓ |  |  |  |
| GSDCA | Bulk | 2011 | ✓ |  |  |  |
| GSNCA | Bulk | 2014 | ✓ |  |  |  |
| MetaGSCA | Bulk | 2021 | ✓ |  |  |  |
| EDDY | Bulk | 2014 | ✓ |  |  |  |
| KEDDY | Bulk | 2019 | ✓ |  |  |  |
| CoGA | Bulk | 2015 | ✓ |  |  |  |
| COTAN | single-cell | 2021 |  | ✓ |  | ✓ |
| CS-CORE | single-cell | 2023 |  | ✓ |  |  |
| GSNCASC R | single-cell | 2025 | ✓ | ✓ |  |  |
| scGSCA | single-cell | 2026 | ✓ | ✓ | ✓ | ✓ |

We focus on three distinct biological comparisons: tumor versus matched adjacent normal tissue in gastric cancer, cancer subtypes within breast tissue, and normal lung versus skin tissue from GTEx. The GTEx comparison serves as a biological positive control. Subject and sample characteristics, cell counts, and analyzed cell types are summarized in Table 2. Each case study addresses two questions. The sample-phenotype test asks whether pathway co-expression differs between phenotypes, and the cell-type interaction test asks whether the magnitude of the co-expression contrast between two cell types differs between phenotypes. For each case study, we describe what the cell-type contrasts add to the sample-phenotype result.

**Table 2.** Overview of the datasets and study designs.

| Case study | Comparison | Study design | No. subjects | No. samples | No. cells | Cell types analyzed | No. pathways |
| --- | --- | --- | --- | --- | --- | --- | --- |
| Gastric cancer | Tumor vs. adjacent normal | Paired | 24 | 48 | 111,140 | Epithelial, fibroblast, myeloid, CD8+ T | 8 |
| Breast cancer | ER+ vs. TNBC | Unpaired | 20 | 20 | 73,730 | Cancer epithelial, CAFs, endothelial, myeloid, T cells | 12 |
| GTEX | Lung vs. skin | Unpaired | 5 | 6 | 40,611 | Epithelial, endothelial, fibroblast, immune | 8 |

### Gastric cancer

The gastric cancer analysis used matched bulk RNA-seq and scRNA-seq data to compare tumor and adjacent normal tissues. The scRNA-seq dataset (HRA002336) contained 111,140 cells and 25,287 genes from 48 samples of 24 patients, including 63,252 tumor cells and 47,888 adjacent normal cells (30). Annotations for 12 major cell types were obtained from the original study: CD8+ T cells, plasma cells, fibroblasts, CD4+ T cells, myeloid cells, B cells, epithelial cells, endothelial cells, mural cells, innate lymphoid cells, mast cells, and glial cells. We visualized the annotated single-cell transcriptomes using UMAP to characterize the major cell populations and their distribution across tumor and adjacent normal samples (Supplementary Figure S1). UMAP visualization showed that the major epithelial, myeloid, B-cell, CD4+ T-cell, CD8+ T-cell, and plasma-cell populations were represented in both normal and tumor samples, while their relative distributions differed between conditions. EPCAM, LYZ, COL1A1, and CD8A expression localized to epithelial, myeloid, fibroblast, and CD8+ T-cell populations, respectively, consistent with the reported cell-type annotations in both normal and tumor samples (Supplementary Figure S2). These data provided the cellular context for the subsequent sample-phenotype and cell-type interaction analyses.

We evaluated eight gastric cancer–related KEGG pathways: Adherens junction (hsa04520) (39), Axon guidance (hsa04360) (40,41), Cell cycle (hsa04110) (42,43), Epithelial cell signaling in *Helicobacter pylori* infection (hsa05120) (44,45), MAPK signaling pathway (hsa04010) (46,47), p53 signaling pathway (hsa04115) (46,48), TGF-beta signaling pathway (hsa04350) (49,50), and Wnt signaling pathway (hsa04310) (51,52). Because tumor and normal observations were obtained from matched samples, we used the paired sample-phenotype test for the scRNA-seq data. Under 200 permutations (limited by runtime; Supplementary Table S1), Axon guidance (*P* = 0.0149) and Wnt signaling (*P* = 0.0199) showed nominally significant phenotype-associated differential co-expression, while other pathways did not reach the nominal significance threshold. As a bulk-level reference, we applied GSNCA to the same eight pathways. Using 500 permutations, seven of the eight showed significant tumor–normal co-expression differences (permutation *P* = 0.0060 − 0.0220), with the strongest signals for TGF-beta signaling (*P* = 0.0060), p53 signaling (*P* = 0.0080), and Cell cycle (*P* = 0.0100). Axon guidance, which was significant in the single-cell sample-phenotype analysis, did not reach the nominal threshold in bulk (*P* = 0.0519). Thus, bulk and single-cell analyses yielded overlapping but distinct patterns of pathway significance.

The cell-type interaction analysis localized one sample-phenotype signal and revealed contrast changes that the pooled test did not detect (500 permutations; unadjusted *P* < 0.05). The Axon guidance signal mapped to the contrast between epithelial and myeloid cells (*P* = 0.0080), whereas Cell cycle in CD8+ T versus myeloid cells (*P* = 0.0379) and TGF-beta signaling in epithelial cells versus fibroblasts (*P* = 0.0319) showed contrast changes without a corresponding sample-phenotype signal. These contrasts involved epithelial, immune, and stromal populations represented in the UMAP and marker-expression visualizations. Whereas those visualizations described the cellular context, the interaction tests identified differences in pathway co-expression contrasts between tumor and adjacent normal tissues. No tested contrast reached nominal significance for Wnt signaling, so its sample-phenotype signal was not resolved into these cell-type contrasts and may reflect changes shared across cell types or differences in cell-type composition. For cell-type-pair pseudo-bulk comparison, five significant pathway-pair signals were detected, none of which overlapped with the cell-type interaction results.

Figures 2 and 3 summarize these results. The sample-phenotype analysis identified differential co-expression in Axon guidance and Wnt signaling, while the cell-type interaction analysis identified phenotype-dependent contrasts involving epithelial, fibroblast, myeloid, and CD8+ T-cell populations. These exploratory findings align with known gastric cancer biology: proliferative programs in gastric adenocarcinoma (31), TGF-beta-dependent stromal and immune remodeling (53,54), Wnt-associated epithelial plasticity (51), and Axon guidance in cell migration and microenvironment remodeling (40). They also match single-cell studies reporting epithelial, fibroblast, myeloid, and T-cell heterogeneity in the gastric tumor microenvironment (55).

**Figure 2.**
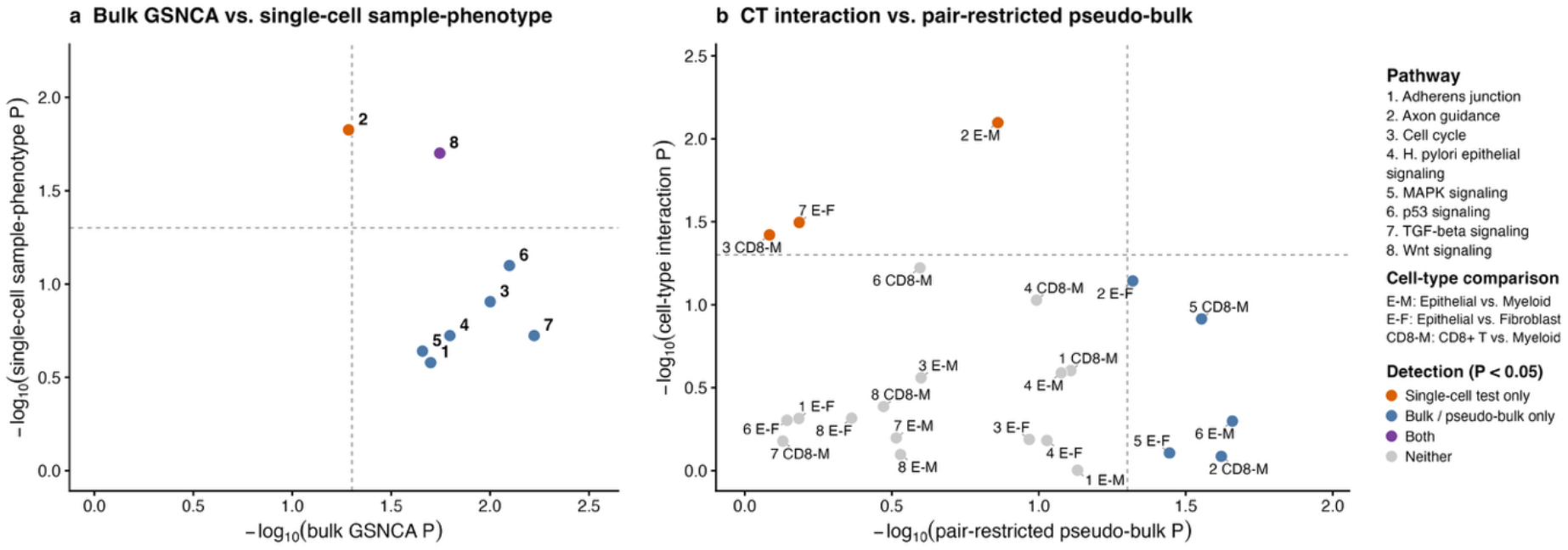
Gastric cancer case study. (a) Comparison of pathway-level *P* values from GSNCA and single-cell sample-phenotype analysis. Each point represents one KEGG pathway. Dashed lines indicate *P* = 0.05. (b) Comparison of cell-type interaction *P* values and cell-type-pair pseudo-bulk GSNCA *P* values for the same pathway and cell-type pair combinations. Each point represents one pathway-pair test. Alt text: Two scatter plots for the gastric cancer case study with dashed lines at *P* = 0.05. Panel (a) plots −*log*10 GSNCA *P* value against −*log*10 single-cell sample-phenotype *P* value for eight KEGG pathways; Axon guidance is significant only in the single-cell test and Wnt signaling in both. Panel (b) plots −*log*10 pseudo-bulk *P* value against −*log*10 cell-type interaction *P* value for pathway and cell-type pair combinations, colored by which test detected them.

**Figure 3.**
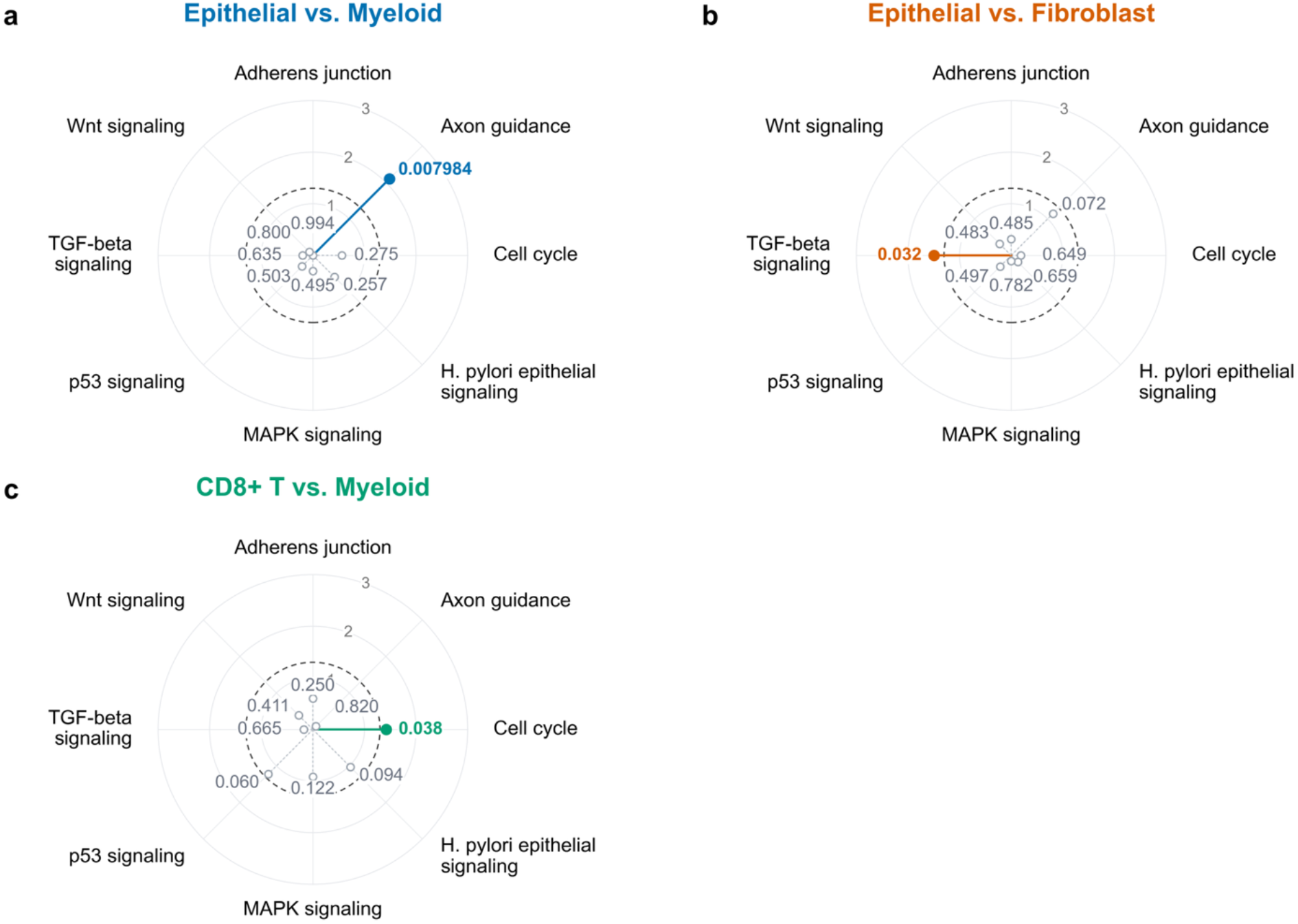
Pathway-level cell-type interaction tests in gastric cancer. Radial plots show cell-type interaction *P* values for eight pathways comparing (a) epithelial and myeloid cells, (b) epithelial cells and fibroblasts, and (c) CD8+ T and myeloid cells. Radial distance represents – log_10,_ *P*, and the dashed circle marks *P* = 0.05. Each spoke represents one of the pathways. Colored filled points with solid spokes indicate *P* < 0.05; gray open points with dotted spokes indicate *P* ≥ 0.05. Labels report unadjusted permutation *P* values based on 500 permutations. Alt text: Three radial plots of cell-type interaction *P* values across eight KEGG pathways in gastric cancer, for epithelial versus myeloid cells, epithelial cells versus fibroblasts, and CD8+ T versus myeloid cells. Spoke length is −*log*10 *P* value and a dashed circle marks *P* = 0.05. One pathway exceeds the threshold in each comparison: Axon guidance (*P* = 0.008), TGF-beta signaling (*P* = 0.032) and Cell cycle (*P* = 0.038), respectively.

### Breast cancer (ER+ versus TNBC)

The breast cancer dataset (GSE176078) includes bulk RNA-seq and scRNA-seq data from primary breast tumors across multiple clinical subtypes (32). We compared ER-positive breast cancer with triple-negative breast cancer (TNBC). The bulk analysis included 11 ER-positive and 9 TNBC samples. The processed scRNA-seq dataset contained 73,730 cells (38,241 ER-positive, 35,489 TNBC). Annotations for 9 major cell types were obtained from the original study: B cells, cancer-associated fibroblasts, cancer epithelial cells, endothelial cells, myeloid cells, normal epithelial cells, plasmablasts, perivascular-like cells, and T cells. UMAP visualization showed the major cell populations entering the sample-phenotype and cell-type interaction analyses (Supplementary Figure S3). CAFs, cancer epithelial cells, myeloid cells, and T cells were present in both subtypes, but at differing proportions, so bulk comparisons cannot separate compositional shifts from within-cell-type co-expression changes. EPCAM, CD3D, LYZ, and COL1A1 expression localized to cancer epithelial, T-cell, myeloid, and CAF populations, respectively, supporting the cell-type annotations in both subtypes (Supplementary Figure S4).

We evaluated twelve Hallmark pathways: angiogenesis (56–58), E2F targets (59,60), epithelial-mesenchymal transition (61,62), estrogen response early (63,64), estrogen response late (65), G2M checkpoint, hypoxia (66–68), IL6-JAK-STAT3 signaling (69–71), interferon gamma response (72,73), MYC targets V1 (74–76), PI3K-AKT-mTOR signaling (77,78), and TNF-*α* signaling via NF-κB (71,79,80). Because the two subtype groups comprised different samples, we applied the unpaired sample-phenotype test. Under 200 permutations, four pathways showed nominally significant subtype-associated differential co-expression: IL6-JAK-STAT3 signaling (*P* = 0.0050), TNF-*α* signaling via NF-κB (*P* = 0.0149), estrogen response late (*P* = 0.0199), and estrogen response early (*P* = 0.0249). As a bulk-level reference, GSNCA was applied to the same 12 pathways using 500 permutations. E2F targets showed a significant ER-positive versus TNBC co-expression difference (*P* = 0.020), but was not significant in the single-cell sample-phenotype analysis (*P* = 0.358). Conversely, the single-cell sample-phenotype analysis detected subtype-associated differences in inflammatory and hormone-response pathways that GSNCA missed: IL6-JAK-STAT3 signaling (bulk *P* = 0.110; single-cell *P* = 0.0050), TNF-*α* signaling via NF-κB (bulk *P* = 0.311; single-cell *P* = 0.0149), estrogen response late (bulk *P* = 0.188; single-cell *P* = 0.0199), and estrogen response early (bulk *P* = 0.385; single-cell *P* = 0.0249).

The cell-type interaction analysis localized the strongest sample-phenotype signal and revealed subtype-dependent contrasts in pathways that the pooled test did not detect (500 permutations; unadjusted *P* < 0.05). IL6-JAK-STAT3 signaling mapped to contrasts between cancer epithelial cells and both immune compartments, whereas angiogenesis, epithelial-mesenchymal transition, G2M checkpoint, and hypoxia showed contrast changes without a corresponding sample-phenotype signal. This included cancer epithelial versus CAF contrasts for angiogenesis (*P* = 0.0020) and epithelial-mesenchymal transition (*P* = 0.0060); cancer epithelial versus myeloid contrasts for G2M checkpoint (*P* = 0.0220) and IL6-JAK-STAT3 signaling (*P* = 0.0279); and cancer epithelial versus T-cell contrasts for hypoxia (*P* = 0.0120) and IL6-JAK-STAT3 signaling (*P* = 0.0439). The estrogen-response and TNF-*α* signals detected by the sample-phenotype test were not resolved into any tested contrast, and no contrast involving endothelial cells reached nominal significance. In the cell-type-pair pseudo-bulk analysis, none of the six combinations identified by the cell-type interaction test reached nominal significance.

Figures 4 and 5 summarize these results. Breast cancer subtype differences are reflected both in pathway-level co-expression changes, particularly inflammatory and hormone-response pathways, and in co-expression contrasts between malignant epithelial cells and immune or stromal compartments. This is consistent with intrinsic subtypes differing in hormone-response and proliferation programs (33,81). The epithelial–immune and epithelial–stromal localization agrees with single-cell and spatial studies describing subtype-associated tumor ecosystems of malignant epithelial cells, T cells, myeloid cells, endothelial cells, and fibroblasts (32).

**Figure 4.**
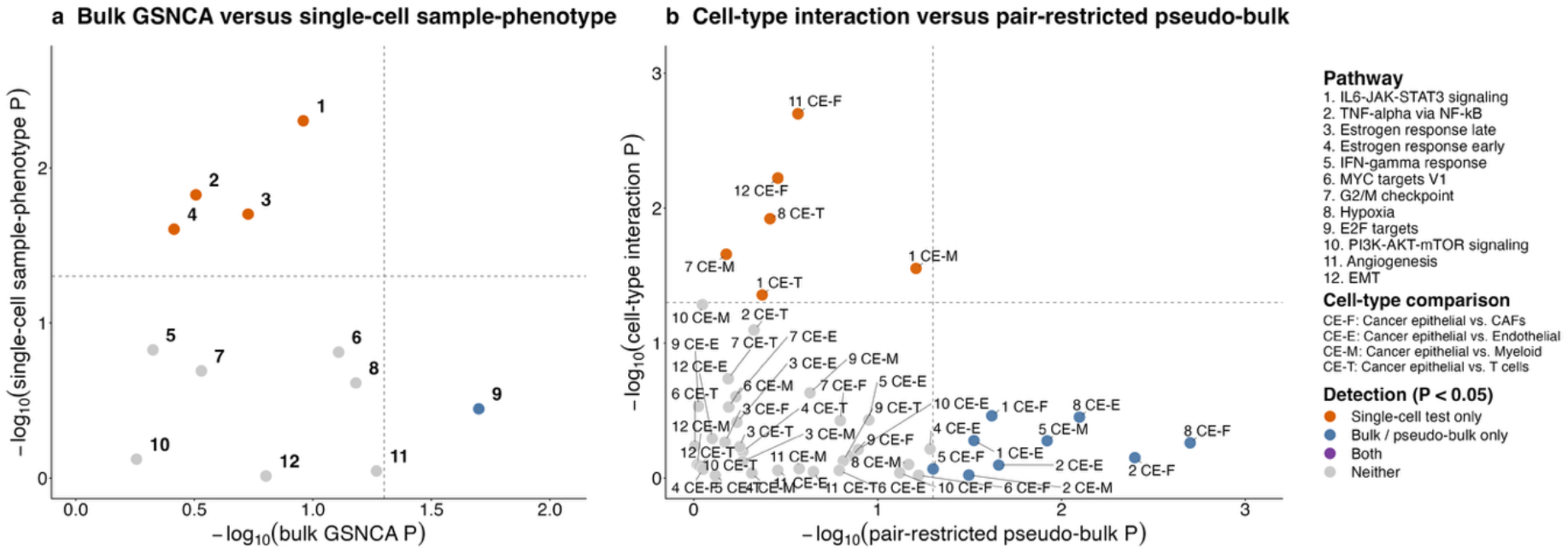
Breast cancer case study. (a) Comparison of pathway-level *P* values from GSNCA and single-cell sample-phenotype analysis. Each point represents one Hallmark pathway. Dashed lines indicate *P* = 0.05. (b) Comparison of cell-type interaction *P* values and cell-type-pair pseudo-bulk GSNCA *P* values for the same pathway and cell-type pair combinations. Each point represents one pathway-pair test. Alt text: Two scatter plots for the breast cancer ER-positive versus TNBC comparison with dashed lines at *P* = 0.05. Panel (a) plots −*log*10 GSNCA *P* value against −*log*10 single-cell sample-phenotype *P* value for 12 Hallmark pathways; IL6-JAK-STAT3, TNF-*α* via NF-κB and both estrogen-response pathways are significant only in the single-cell test, and E2F targets only in bulk. Panel (b) compares cell-type interaction and pseudo-bulk *P* values for pathway and cell-type pair combinations.

**Figure 5.**
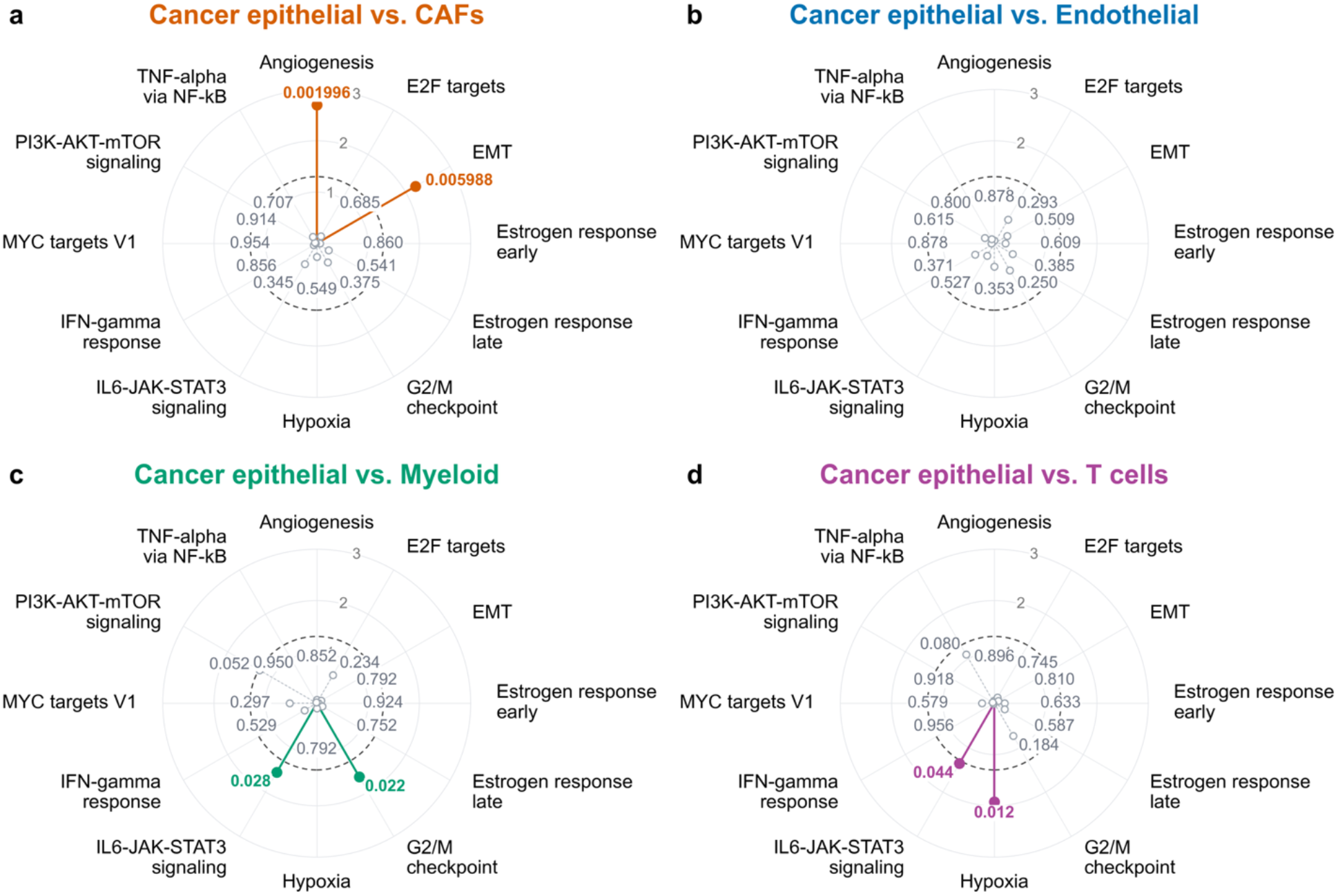
Pathway-level cell-type interaction tests in breast cancer ER+ and TNBC subtypes. Radial plots show cell-type interaction *P* values for 12 pathways comparing cancer epithelial cells with (a) cancer-associated fibroblasts (CAFs), (b) endothelial cells, (c) myeloid cells, and (d) T cells across ER+ and TNBC samples. Labels report unadjusted permutation *P* values based on 500 permutations. Radial distance represents – log_10,_ *P*, and the dashed circle marks *P* = 0.05. Colored filled points with solid spokes indicate *P* < 0.05; gray open points with dotted spokes indicate *P* ≥ 0.05. Alt text: Four radial plots of cell-type interaction *P* values across 12 Hallmark pathways, comparing cancer epithelial cells with CAFs, endothelial cells, myeloid cells and T cells in ER-positive versus TNBC tumors. Spoke length is −*log*10 *P* value and a dashed circle mark *P* = 0.05. Significant pathways are angiogenesis and epithelial-mesenchymal transition for CAFs, none for endothelial cells, IL6-JAK-STAT3 and G2M checkpoint for myeloid cells, and IL6-JAK-STAT3 and hypoxia for T cells.

### GTEx lung versus skin

To evaluate whether the scGSCA framework could recover expected tissue-specific co-expression differences outside the tumor setting, we compared GTEx lung and skin tissues as a biological positive control using matched bulk RNA-seq and scRNA-seq resources. Because these tissues differ in cellular composition and function, we expected widespread co-expression differences across pathways. The GTEx database (34) includes bulk RNA-seq and scRNA-seq data for skin and lung. The bulk analysis used 3 lung and 3 skin samples. The scRNA-seq dataset contained 40,611 cells (35,284 lung, 5,327 skin). Annotations for 10 major cell types were obtained from previous studies: epithelial cells, immune cells, endothelial cells, fibroblasts, pericytes/smooth muscle cells, adipocytes, melanocytes, sweat gland cells, sebaceous gland cells, and unknown cells. UMAP visualization showed the major cell populations entering the sample-phenotype and cell-type interaction analyses (Supplementary Figure S5). Epithelial, fibroblast, endothelial, and immune cells were present in both tissues, but at differing proportions, so bulk comparisons cannot separate compositional shifts from within-cell-type co-expression changes. EPCAM, PECAM1, COL1A1, and PTPRC expression localized to epithelial, endothelial, fibroblast, and immune cell populations, respectively, supporting the cell-type annotations in both tissues (Supplementary Figure S6).

We evaluated eight Hallmark pathways: apoptosis (82–84), complement (85–87), epithelial-mesenchymal transition (88–90), inflammatory response (91–93), interferon gamma response (94,95), Notch signaling (96,97), TGF-beta signaling (98,99), and Wnt/beta-catenin signaling (100,101). In the scRNA-seq sample-phenotype analysis, seven of the eight pathways showed significant tissue-associated co-expression remodeling under 200 permutations (*P* = 0.005 to 0.0448), with the strongest signals for epithelial-mesenchymal transition and TGF-beta signaling. Notch signaling showed a marginal trend (*P* = 0.0647). As a bulk-level reference, GSNCA identified all eight pathways with significant lung– skin co-expression differences (all *P* = 0.0020).

The cell-type interaction analysis added information on where and in which direction these tissue differences arise. Under 500 permutations, nominally significant contrasts spanned all eight pathways, including Notch signaling, which was only marginal in the sample-phenotype test, and were concentrated in endothelial cell versus immune, epithelial cell versus endothelial cell, and fibroblast versus immune comparisons. Of these, 26 attained the minimum attainable permutation *P* value of 1/501 (approximately 0.001996), reflecting the finite resolution of the permutation procedure rather than identical effect sizes. In every significant contrast, *D* was larger in lung than in skin, indicating stronger cell-type contrast statistics in lung than in skin. The cell-type-pair pseudo-bulk analysis identified only two significant pathway-pair signals: apoptosis for endothelial versus immune cells and complement for epithelial versus immune cells (both *P* = 0.0160). Figures 6 and 7 summarize these results. The lung–skin comparison demonstrated tissue-associated differential co-expression across multiple pathways and tissue-dependent co-expression contrasts involving epithelial, endothelial, fibroblast, and immune populations. The cell-type interaction results complemented the bulk and pseudo-bulk analyses by characterizing contrasts between cell populations rather than co-expression in pooled profiles.

**Figure 6.**
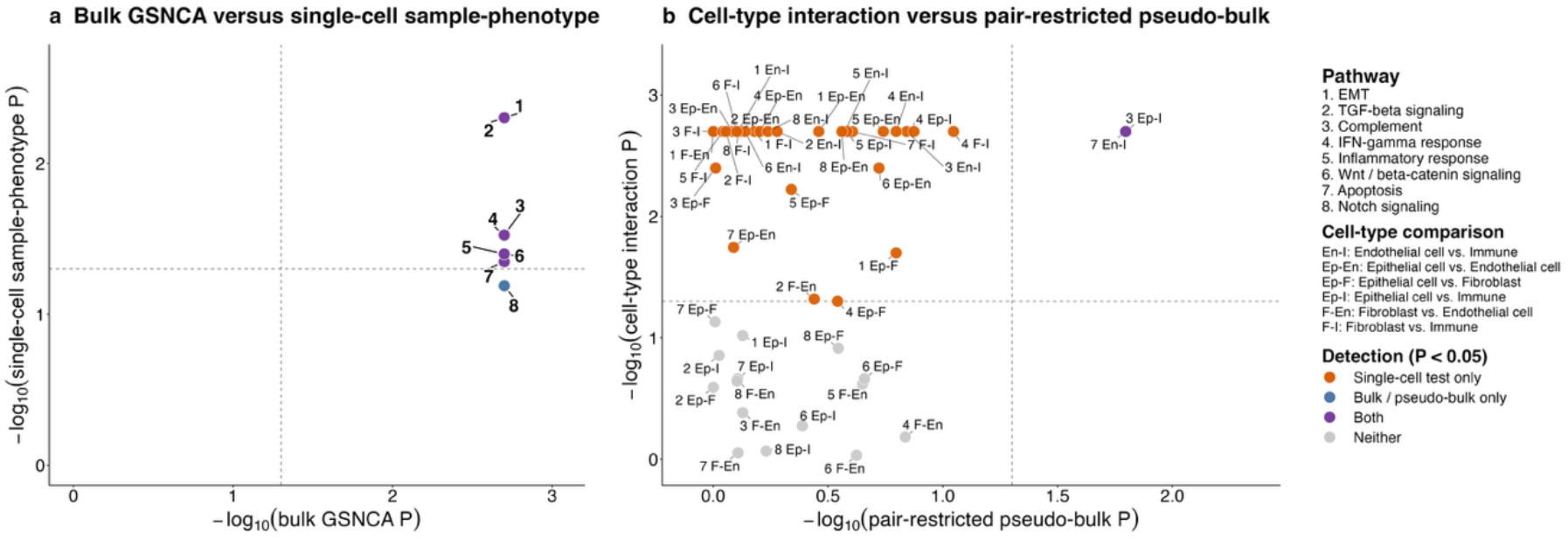
GTEx lung versus skin case study. (a) Comparison of pathway-level *P* values from GSNCA and single-cell sample-phenotype analysis. Each point represents one Hallmark pathway. Dashed lines indicate *P* = 0.05. (b) Comparison of cell-type interaction *P* values and cell-type-pair pseudo-bulk GSNCA *P* values for the same pathway and cell-type pair combinations. Each point represents one pathway-pair test. Alt text: Two scatter plots for the GTEx lung versus skin comparison with dashed lines at *P* = 0.05. Panel (a) shows that all eight Hallmark pathways are significant in GSNCA and seven are also significant in the single-cell sample-phenotype test, with Notch signaling significant only in bulk. Panel (b) shows that most pathway and cell-type pair combinations are significant only in the cell-type interaction test; two are significant in both tests.

**Figure 7.**
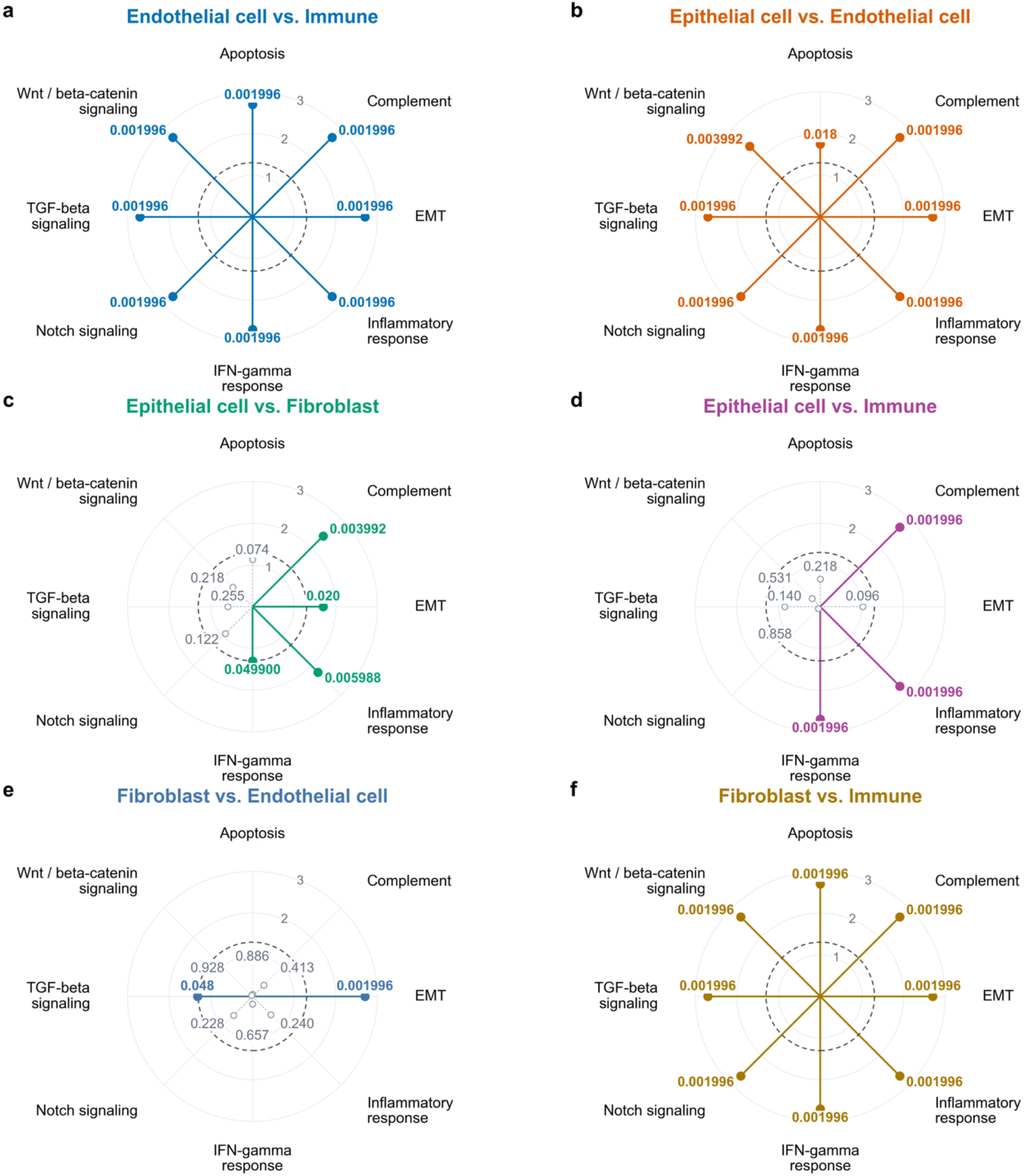
Pathway-level cell-type interaction tests in GTEx lung and skin tissues. Radial plots show cell-type interaction *P* values for eight pathways comparing (a) endothelial and immune cells, (b) epithelial and endothelial cells, (c) epithelial cells and fibroblasts, (d) epithelial and immune cells, (e) fibroblasts and endothelial cells, and (f) fibroblasts and immune cells. Radial distance represents – log_10,_ *P*, and the dashed circle marks *P* = 0.05. Colored filled points with solid spokes indicate *P* < 0.05; gray open points with dotted spokes indicate *P* ≥ 0.05. Labels report unadjusted permutation *P* values based on 500 permutations. Alt text: Six radial plots of cell-type interaction *P* values across eight Hallmark pathways in GTEx lung versus skin, for pairs of epithelial, endothelial, fibroblast and immune cells. Spoke length is – log_10,_ *P* and a dashed circle marks *P* = 0.05. All eight pathways are significant for endothelial versus immune, epithelial versus endothelial, and fibroblast versus immune cells, most at the minimum attainable P of 0.001996. Fewer pathways are significant for the other three pairs.

Runtime was recorded separately for sample-phenotype and cell-type interaction analyses. Hardware specifications and runtimes are provided in Supplementary Table S1.

## Discussion

In this work, we developed scGSCA, a gene-set-level framework for detecting differential co-expression patterns in scRNA-seq data. By modeling gene expression with ZINBMM and applying residual-based gene-set co-expression analysis, scGSCA evaluates coordinated changes in pathway structure while accounting for overdispersion, excess zeros, and subject-level dependence. The framework includes two complementary tests: the sample-phenotype analysis, which detects sample-phenotype-associated pathway co-expression changes after adjusting for cell-type heterogeneity, and the cell-type interaction analysis, which identifies sample-phenotype-dependent changes in cell-type-specific pathway contrasts. Together, these tests enable investigators to examine pathway co-expression at both the population and cell-type-contrast levels within a single subject-aware framework.

Across three case studies, scGSCA identified pathway co-expression changes that bulk analysis did not fully capture. The sample-phenotype test detected signals absent from GSNCA, such as Axon guidance in gastric cancer and inflammatory and estrogen-response pathways in breast cancer. The cell-type interaction test localized phenotype-dependent contrasts, most of which involved epithelial cells and stromal or immune populations. In the GTEx positive control, scGSCA recovered the expected tissue differences and localized them to cell-type contrasts that pseudo-bulk analysis largely missed.

Bulk RNA-seq and scRNA-seq can differ not only in statistical power, but also in biological interpretation, because they capture tissue-averaged versus cell-resolved expression signals. scGSCA explicitly accounts for cell-type effects and within-subject dependence, whereas bulk analysis evaluates co-expression in aggregated tissue profiles. Similarly, limited overlap between the cell-type interaction test and the cell-type-pair pseudo-bulk analysis is expected because the two address different questions: the interaction test asks whether the co-expression contrast between two cell types differs across phenotypes, whereas the pseudo-bulk analysis asks whether pathway co-expression differs between phenotypes after the two cell populations are pooled.

As a complementary analysis, we applied GSNCASCR to the same gastric cancer KEGG pathways and breast cancer Hallmark pathways analyzed by scGSCA (Supplementary Table S2). GSNCASCR uses CS-CORE to estimate cell-type-specific gene-gene correlations from raw UMI counts while accounting for measurement noise. It supports GSNCA-based pathway comparisons and gene-level network connectivity summaries. In our analysis, we focused on gene-level connectivity changes and obtained gene-by-cell-type matrices of network weight difference scores (Supplementary Data S2), without performing pathway-level significance testing. In gastric cancer, MAPK signaling and Cell cycle had the largest median scores among the eight pathways. In breast cancer, E2F targets, MYC targets V1, and G2M checkpoint had the largest median scores. These descriptive scores were not normalized for gene-set size and should not be interpreted as pathway-level significance or directly comparable effect sizes. This comparison illustrates differences in analytical outputs but does not evaluate the relative power or false-positive control of the two methods. scGSCA additionally models subject-level dependence through random effects and implements permutation procedures that preserve subject membership. It also provides a cell-type interaction test to assess whether co-expression contrasts between cell types differ across phenotypes.

A key feature of scGSCA is that co-expression is estimated from model-based residuals rather than raw counts, whose correlations are distorted by zero inflation, overdispersion, library-size variation, and repeated measurement within subjects. The fixed-effect structure can include biological or technical covariates relevant to the research question. Subject-level random effects and subject-preserving permutation guard against the Type I error inflation that arises when cells are treated as independent, without requiring closed-form null distributions.

We further examined whether the cell-type interaction *P* values were stable to the number of permutations by running representatives of significant, borderline, and non-significant tests with increasing permutation numbers. The resulting *P* values remained qualitatively stable, suggesting that the reported cell-type interaction findings were not driven by Monte Carlo variability from a small number of permutations (Supplementary Figure S7).

As described in Materials and Methods, permutation validity requires exchangeability under the null within the permitted rearrangements, and dependence among observations must be preserved (102). We evaluated the joint permutation procedure based on *δ* in four null simulation settings, each consisting of 500 datasets with 200 permutations per dataset. Rejection rates were evaluated at the nominal *α* = 0.05 level. The four null simulation settings assessed rejection behavior under distinct latent correlation structures with matched absolute loading patterns, baseline calibration with identical cell types and no subject effects, the influence of subject-level dependence with identical cell types, and the influence of cell-type heterogeneity without subject effects. The Type I error rate in each setting was 0.014 (CI: [0.0056, 0.0286]), 0.062 (CI: [0.0425, 0.0869]), 0.042 (CI: [0.0262, 0.0635]) and 0.006 (CI: [0.0012, 0.0174]), respectively (Supplementary Figures S8 and S9). These results indicate setting-dependent calibration, with conservative behavior when cell types differed in the absence of subject effects.

Several limitations remain. First, fitting gene-wise mixed models and performing permutations is computationally intensive, particularly for sample-phenotype analyses. Nevertheless, all analyses reported here were completed on standard laptops; multi-core or cluster computing can reduce runtime for large pathway collections or many cell-type contrasts. The number of permutations limits both the minimum attainable *P* value, 1/(*n*_*perm*_ + 1), and the Monte Carlo precision of the permutation estimate.

Separately, when many pathways or cell-type contrasts are tested, multiplicity should be controlled, and *P* values can be adjusted using standard false discovery rate procedures such as the Benjamini–Hochberg method. These case studies illustrate the method’s application rather than establish definitive pathway associations; we therefore report unadjusted *P* values and interpret the findings as exploratory and hypothesis-generating. The number of subject-level assignments limits permutation resolution for the sample-phenotype test; the interaction test additionally includes within-subject cell-type rearrangements. With few subjects per group, only a small number of distinct label assignments exist, and the attainable *P* values are limited regardless of how many permutations are requested. This is most relevant to the GTEx comparison, which included five subjects.

Second, like all pathway-based methods, performance depends on the accuracy of predefined gene sets, and results are additionally conditional on cell-type annotations imported from prior studies rather than derived here. Third, Type I error control for the cell-type interaction test is setting-dependent: empirical rejection rates ranged from 0.006 to 0.062 across the four null settings, indicating mild anti-conservatism when cell types are identical and subject effects are absent, and conservative behavior when cell types differ. The interaction procedure requires within-subject exchangeability of cell-type labels, an assumption stronger than equality of the between-cell-type co-expression distances across phenotypes.

The reported simulations assess calibration in selected settings but do not establish validity under the general interaction null. Fourth, the current implementation is restricted to pairwise group comparisons and selected cell-type contrasts. Future extensions to multi-condition designs, continuous covariates, and complex interactions would broaden its scope. Because the test statistic compares leading eigenvectors of absolute residual correlation matrices, it summarizes the dominant axis of co-expression and does not distinguish gains from losses of coordination, so interpreting the direction of remodeling requires inspecting the underlying correlation structures. In addition, scGSCA may be extendable to spatial transcriptomic data when cell- or spot-level expression profiles and cell-type annotations are available. Finally, further evaluation in larger, independent cohorts would help establish generalizability.

These constraints bound the scope of the present results rather than the approach itself. scGSCA offers a model-based and interpretable framework for single-cell gene-set co-expression analysis. By combining model-based residualization with gene-set-level testing, scGSCA characterizes phenotype-associated differential co-expression and changes in co-expression contrasts between cell types, complementing conventional differential expression analysis. Its use of count matrices and predefined gene sets makes it broadly applicable to multi-subject single-cell studies.

## Supplementary data

Supplementary data is available at NAR online.

## Acknowledgements

The authors thank the investigators and participants of the original gastric cancer, breast cancer, and GTEx studies for making these data available. We also thank the developers of the public databases and software packages used in this work. The generative AI tool ChatGPT (version 5.6-sol; OpenAI) was used in the preparation of this work as follows: (i) to create the illustrative graphical abstract and the schematic in Figure 1, which depict the scGSCA workflow and do not present data or results; (ii) to translate portions of the original R implementation into C++ for the Rcpp-based components of the scGSCA package; and (iii) to check grammar and spelling and to shorten selected paragraphs of the manuscript prior to submission. All AI-generated content was reviewed and edited by the authors; the C++ code was verified against the original R implementation. The authors take full responsibility for the content of this article and the accompanying software.

## Author contributions

Zhuoli Jin: Methodology, Software, Data curation, Formal analysis, Validation, Visualization, Writing – original draft, Writing – review & editing. Rebecca Irlmeier: Methodology, Software, Data curation, Formal analysis, Validation, Visualization, Writing – original draft, Writing – review & editing. Jie Ping: Data curation, Software, Formal analysis, Writing – original draft. Yuan Wang: Methodology, Writing – review & editing. Yan Guo: Conceptualization, Data curation, Supervision, Writing – review & editing. Fei Ye: Conceptualization, Methodology, Supervision, Funding acquisition, Writing – original draft, Writing – review & editing.

## Conflict of Interest

The authors declare no conflict of interest.

## Funding

This work was partially supported by the National Institutes of Health: The National Cancer Institute [P30CA240139].

## Ethics Statement

This study involved secondary analysis of previously published, de-identified human sequencing data obtained from public repositories (GSA-Human, GEO, the GTEx Portal, and dbGaP). Ethical approval and informed consent were obtained by the original studies, as described in their respective publications. No new human participants were recruited, and no identifiable information was accessed for this work. All data were accessed and used in accordance with the data use policies of the respective repositories.

## Data Availability

The datasets analyzed in this study are publicly available. The gastric cancer scRNA-seq data are available from the Genome Sequence Archive for Human (GSA-Human) at the National Genomics Data Center under accession HRA002336, and the matched bulk RNA-seq data are available from the Gene Expression Omnibus (GEO) under accession GSE206785. The breast cancer bulk RNA-seq and scRNA-seq data are available from GEO under accession GSE176078. The GTEx bulk RNA-seq and scRNA-seq data are available through the GTEx Portal (https://gtexportal.org) and the database of Genotypes and Phenotypes (dbGaP) under accession phs000424. KEGG pathway gene sets were obtained from the KEGG database and Hallmark gene sets from the Molecular Signatures Database (MSigDB). The scGSCA R package and the analysis scripts used in this study are freely available at https://github.com/Ye-Lab-UMiami/scGSCA. The version used in this study is archived at Zenodo (https://doi.org/10.5281/zenodo.23144513).

## Notes

https://github.com/Ye-Lab-UMiami/scGSCA

https://doi.org/10.5281/zenodo.23144513

### Competing Interest Statement

The authors have declared no competing interest.

