## Supplement Figure for "scGSCA: gene-set differential co-expression analysis for single-cell RNA sequencing data"

### Figures

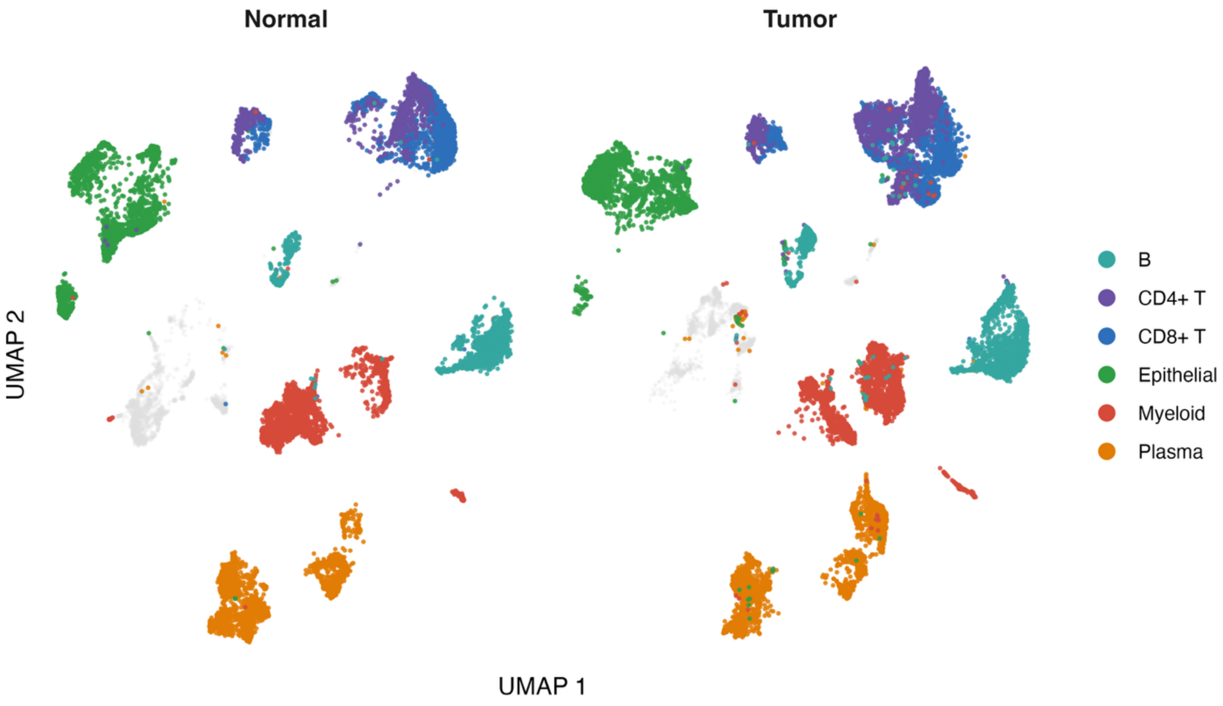


**Figure S1. UMAP visualization of annotated cell populations in the gastric cancer scRNA-seq dataset.** Cells are split by adjacent normal and tumor tissue and colored by major cell type (epithelial, myeloid, CD8+ T, CD4+ T, B, plasma); other types are gray.

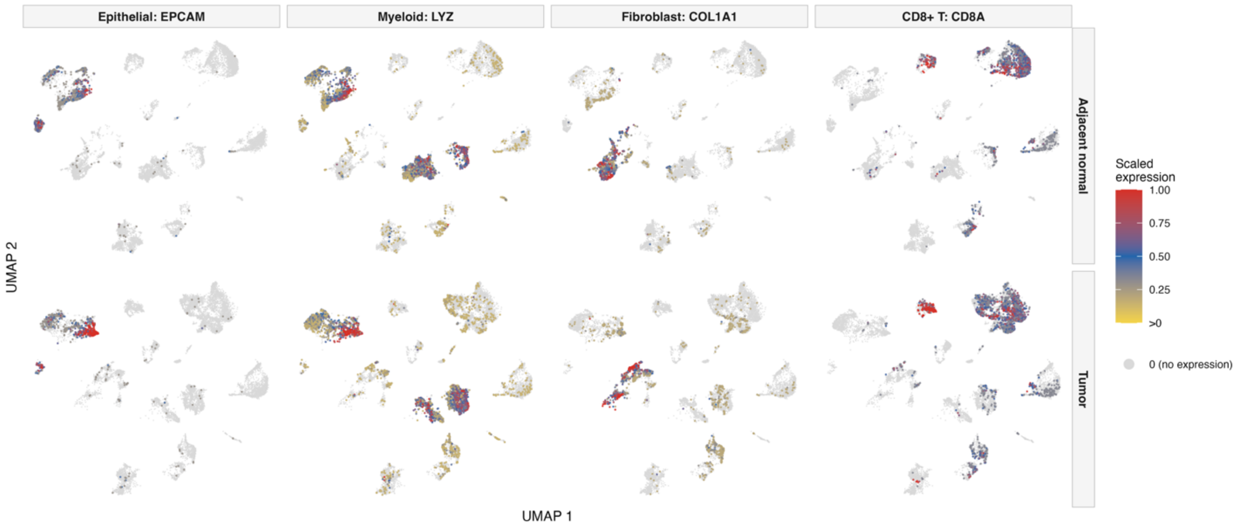

**Figure S2. Marker-specific UMAP visualization of the gastric cancer scRNA-seq dataset.** Cells are split by adjacent normal and tumor tissue and colored by scaled expression of canonical markers (EPCAM, epithelial; LYZ, myeloid; COL1A1, fibroblast; CD8A, CD8+ T).

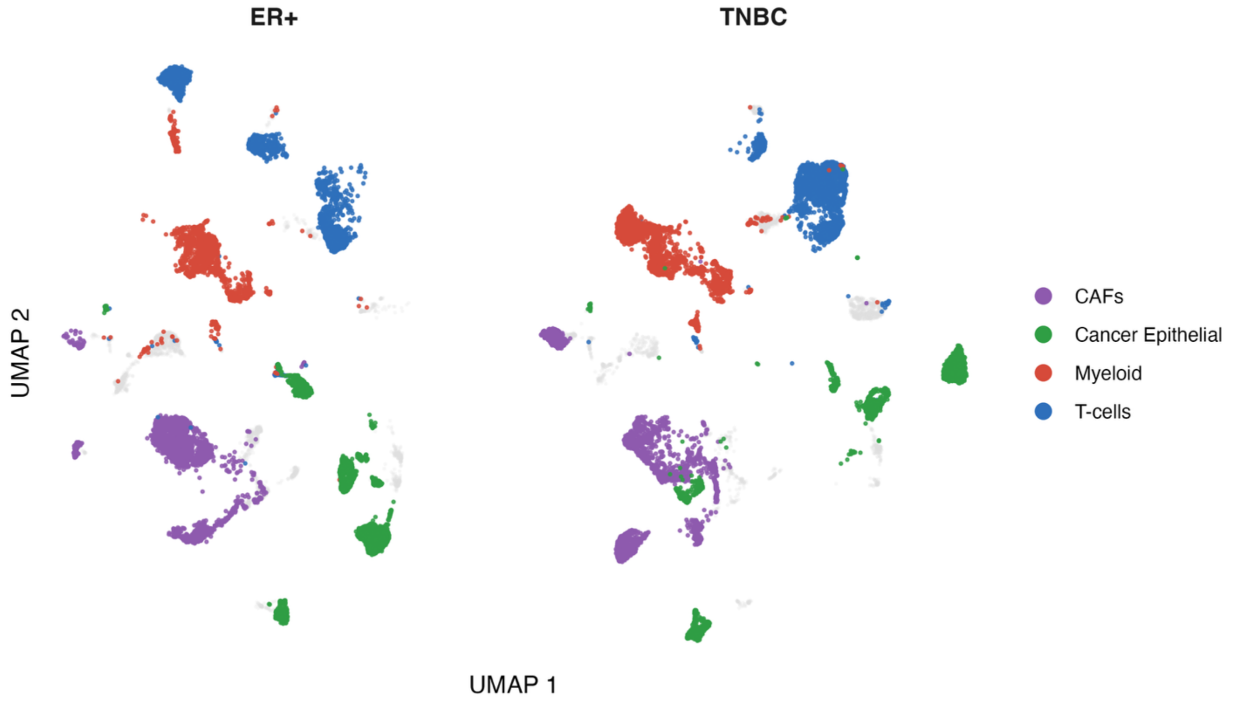


**Figure S3. UMAP visualization of annotated cell populations in the breast cancer scRNA-seq dataset.** Cells are split by ER-positive and TNBC samples. Cells are colored by major cell type (cancer epithelial, myeloid, T cell, CAF); other types are gray.

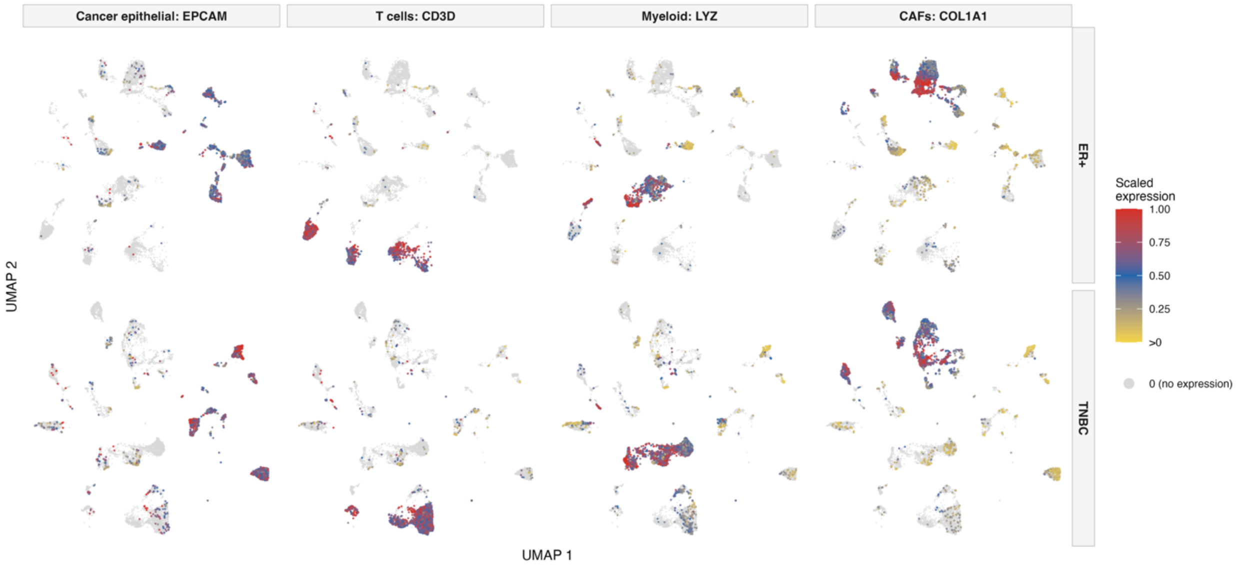

**Figure S4. Marker-specific UMAP visualization of the breast cancer scRNA-seq dataset.** Cells are split by ER-positive and TNBC samples and colored by scaled expression of canonical markers (EPCAM, cancer epithelial; CD3D, T cell; LYZ, myeloid; COL1A1, CAF).

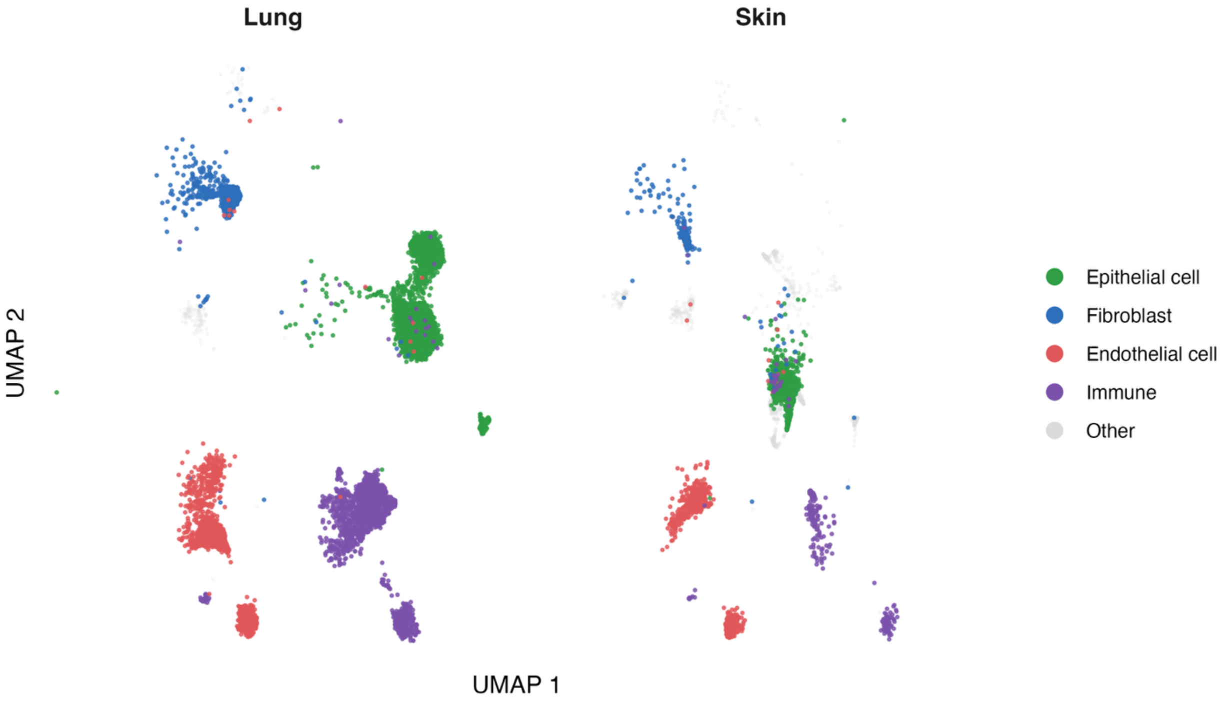


**Figure S5. UMAP visualization of annotated cell populations in the GTEx scRNA-seq dataset.** Cells are split by lung and skin tissue and colored by major cell type (epithelial, endothelial, fibroblast, immune); other types are gray.

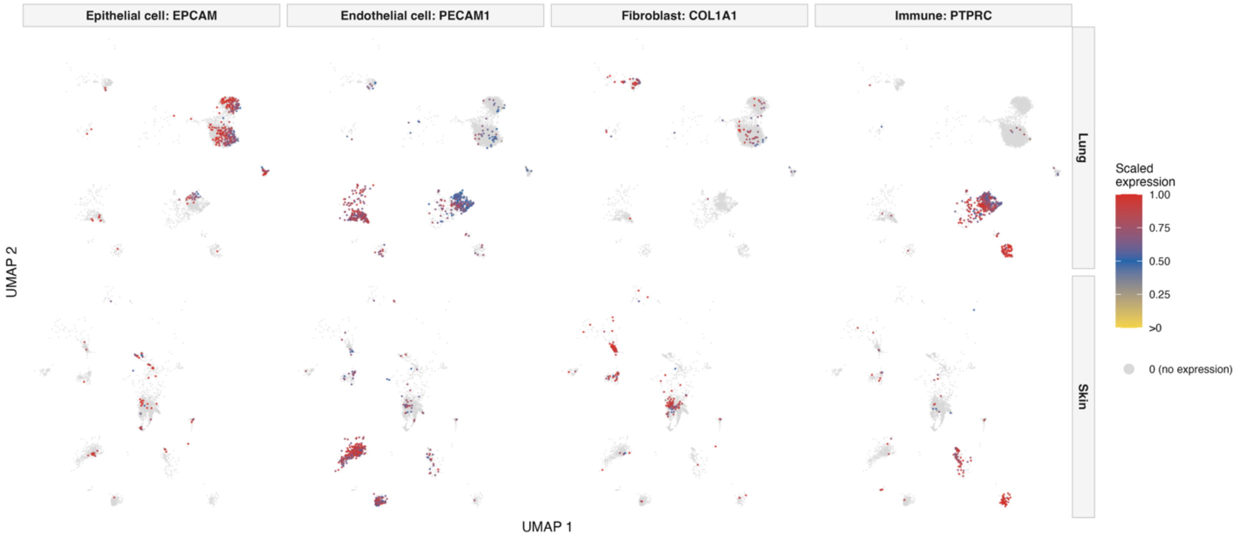

**Figure S6. Marker-specific UMAP visualization of the GTEx scRNA-seq dataset.** Cells are split by lung and skin tissue and colored by scaled expression of canonical markers (EPCAM, epithelial; PECAM1, endothelial; COL1A1, fibroblast; PTPRC, immune).

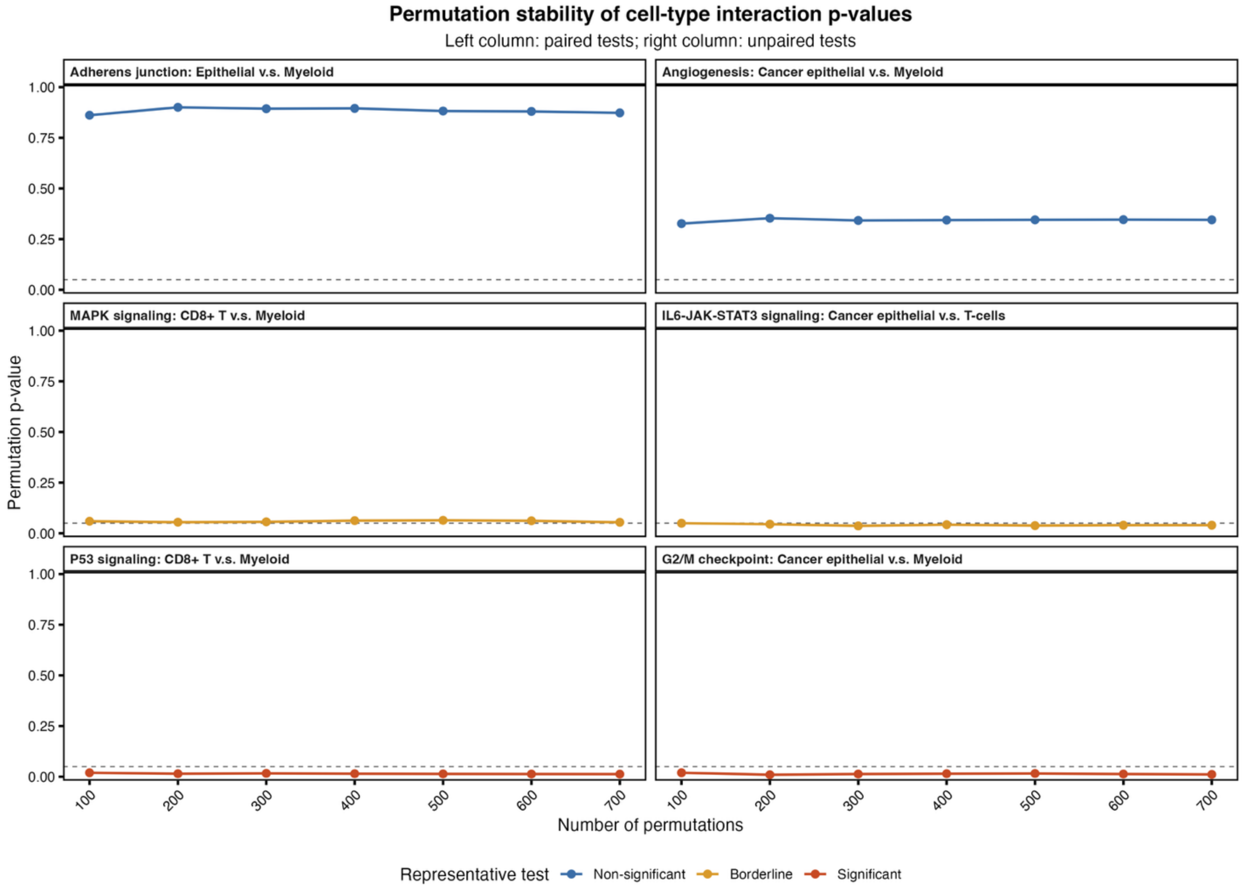


**Figure S7. Cell-type interaction *P* values.** Representative paired and unpaired cell-type-interaction tests using increasing numbers of permutations. Each panel shows the permutation $P$ value of $\delta$ for one pathway and cell-type pair. The dashed horizontal line indicates $P = 0.05$. Non-significant, borderline, and significant examples are shown for both paired and unpaired designs.

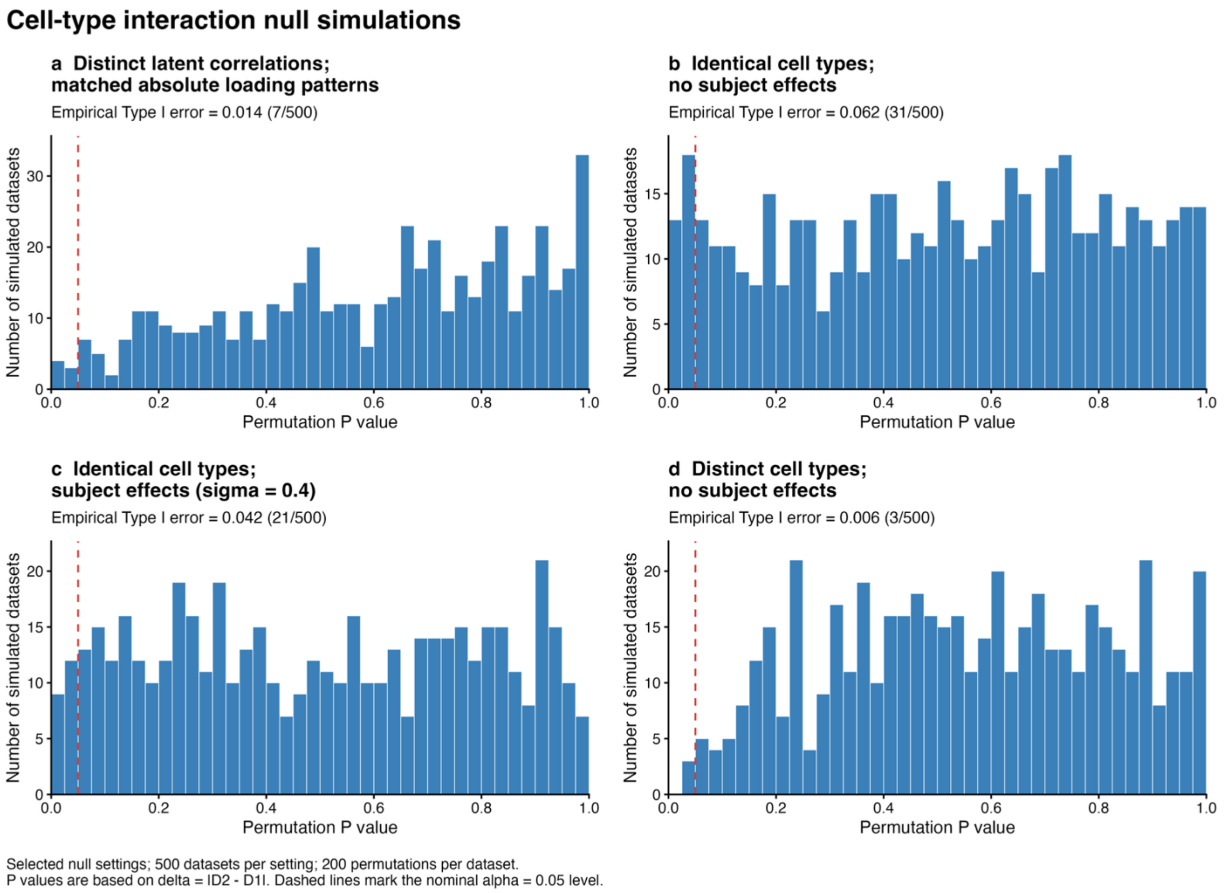


**Figure S8. Cell-type interaction Type I error simulation**. Distribution of permutation $P$ values based on $\delta = |D_2 - D_1|$ under four null settings: (a) distinct cell types with latent correlation matrices related by gene-wise sign reversals, giving matched absolute loading patterns; (b) identical cell types without subject effects; (c) identical cell types with subject effects ($\sigma = 0.4$); (d) distinct cell types without subject effects. Each setting comprises 500 simulated datasets with 200 permutations per dataset, and dashed lines mark the nominal $\alpha = 0.05$ level.

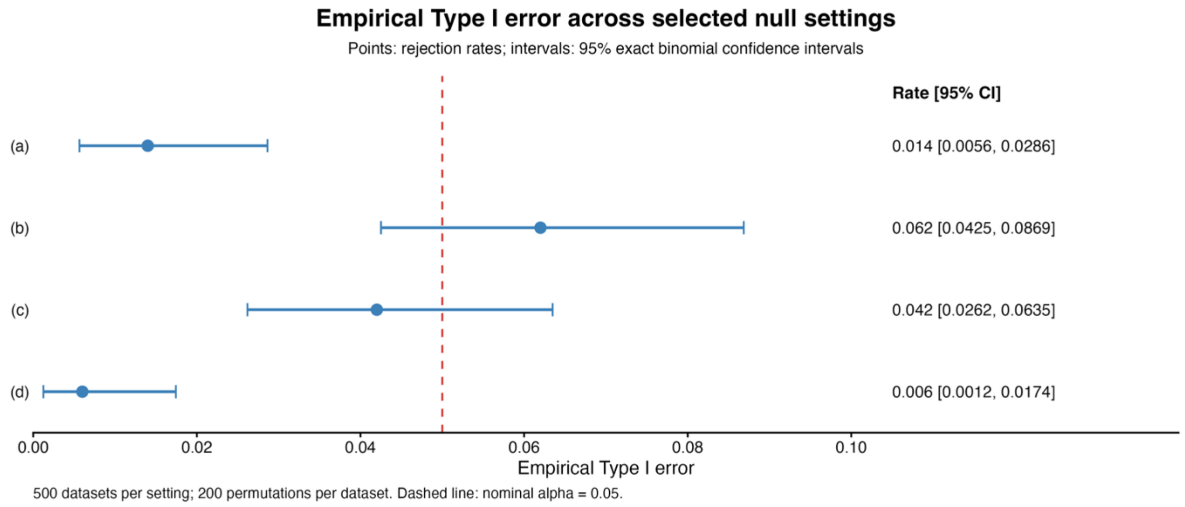

**Figure S9. Empirical Type I error rates across four null simulation settings.** Points show empirical rejection rates at the nominal $\alpha = 0.05$ level, and horizontal bars indicate 95% exact binomial confidence intervals. Settings (a) to (d) correspond to those in Figure S8. The red dashed line marks the nominal rejection rate of 0.05.
