## Supplement Table for "scGSCA: gene-set differential co-expression analysis for single-cell RNA sequencing data"

### Tables

Computational time varied across datasets and analysis types. Using 4-8 CPU cores, cell-type interaction tests averaged $<$ 1 hour per pathway-cell-type contrast in GTEx (~41,000 cells) and $<$ 3 hours in the cancer datasets (~74,000-111,000 cells). Sample-phenotype tests were more computationally demanding, averaging approximately 10 hours per pathway in GTEx and $>$ 10 hours in both cancer datasets.

**Table S1. Analysis settings for the scGSCA case studies.** Numbers of samples, cells, pathways, cell-type contrasts and permutations are listed for the sample-phenotype and cell-type interaction analyses in each case study. All analyses were performed using 4-8 CPU cores. Configuration A: MacBook Pro, Apple M4 Pro, 14-core CPU 10 performance + 4 efficiency cores, 48 GB memory. Configuration B: MacBook Pro, Apple M5, 10-core CPU 4 performance + 6 efficiency cores, 16 GB memory.

| Case Study | Analysis | No. Samples | No. Cells | No. Pathways | No. Cell-type Contrasts | No. Permutations |
|---|---|---|---|---|---|---|
| Gastric Cancer | sample-phenotype | 48 | 111140 | 8 | — | 200 |
| Gastric Cancer | cell-type interaction | 48 | 111140 | 8 | 3 | 500 |
| Breast Cancer | sample-phenotype | 20 | 73730 | 12 | — | 200 |
| Breast Cancer | cell-type interaction | 20 | 73730 | 12 | 4 | 500 |
| GTEx | sample-phenotype | 6 | 40611 | 8 | — | 200 |
| GTEx | cell-type interaction | 6 | 40611 | 8 | 6 | 500 |

**Table S2. Summary of GSNCASCR network weight difference scores in gastric and breast cancer.** Analyses compared (A) tumor and adjacent normal tissues across eight KEGG pathways in gastric cancer and (B) ER-positive and TNBC tumors across 12 Hallmark gene sets in breast cancer. Each score sums absolute between-group differences in gene–gene co-expression estimates involving a given gene. Differences below 0.2 were set to zero, and edge-level values were rounded to two decimal places before summation.

**A. Gastric cancer**

| KEGG pathway | KEGG ID | No. genes | No. cell types | Median difference score (Q1–Q3) |
|---|---|---|---|---|
| p53 signaling | hsa04115 | 74 | 12 | 12.03 (5.27–19.64) |
| Wnt signaling | hsa04310 | 170 | 12 | 26.96 (0.00–45.64) |

| | | | | |
|---|---|---|---|---|
| TGF-beta signaling | hsa04350 | 107 | 12 | 17.14 (0.64–28.02) |
| Axon guidance | hsa04360 | 177 | 12 | 27.88 (0.00–45.11) |
| MAPK signaling | hsa04010 | 282 | 12 | 46.27 (9.06–76.58) |
| Cell cycle | hsa04110 | 154 | 12 | 35.88 (21.50–52.70) |
| Adherens junction | hsa04520 | 86 | 12 | 16.77 (9.00–26.11) |
| Epithelial cell signaling in Helicobacter pylori infection | hsa05120 | 70 | 12 | 13.69 (4.70–21.04) |

**B. Breast cancer**

| **Hallmark gene set** | **No. genes** | **No. cell types** | **Median difference score (Q1–Q3)** |
|---|---|---|---|
| Estrogen Response Early | 194 | 9 | 35.98 (22.80–58.26) |
| Estrogen Response Late | 194 | 9 | 33.92 (19.64–55.13) |
| Epithelial Mesenchymal Transition | 198 | 9 | 29.82 (15.24–46.99) |
| Interferon Gamma Response | 196 | 9 | 33.29 (19.93–51.21) |
| TNFA Signaling via NFKB | 198 | 9 | 18.50 (10.37–32.87) |
| IL6-JAK-STAT3 Signaling | 86 | 9 | 12.09 (5.74–20.73) |
| Hypoxia | 193 | 9 | 29.18 (16.59–46.83) |
| Angiogenesis | 36 | 9 | 3.92 (2.01–6.96) |
| G2M Checkpoint | 189 | 9 | 47.04 (28.37–69.71) |
| E2F Targets | 195 | 9 | 55.02 (36.64–77.09) |
| MYC Targets V1 | 193 | 9 | 54.26 (38.04–74.51) |
| PI3K-AKT-MTOR Signaling | 103 | 9 | 26.67 (17.49–37.91) |
